# Identification of QTL-by-Environment Interaction by Controlling Polygenic Background Effect

**DOI:** 10.1101/2024.12.29.630685

**Authors:** Fuping Zhao, Lixian Wang, Shizhong Xu

**Affiliations:** State Key Laboratory of Animal Biotech Breeding, Institute of Animal Science, Chinese Academy of Agricultural Sciences, Beijing 100193, China; Department of Botany and Plant Sciences, University of California, Riverside, CA 92521, USA

**Keywords:** Q×E interaction, Main effect, Meta-analysis, Mixed model, Polygenic background

## Abstract

The QTL by environment interaction (Q×E) effect is hard to detect because there are no effective ways to control the genomic background. In this study, we propose a novel linear mixed model that simultaneously analyzes data from multiple environments to detect Q×E interactions. This model incorporates two different kinship matrices derived from the genome-wide markers to control both main and interaction polygenic background effects. Simulation studies demonstrated that our approach was more powerful than the meta-analysis and inclusive composite interval mapping methods. We further analyzed four agronomic traits of rice across four environments. A main effect QTL was identified for 1000-grain weight (KGW), while no QTLs were found for tiller number. Additionally, a large QTL with a significant Q×E interaction was detected on chromosome 7 affecting grain number, yield and KGW. This region harbored two important genes, *PROG1* and *Ghd7*. Furthermore, we applied our mixed model to analyze lodging in barley across six environments. The six regions exhibiting Q×E interaction effects identified by our approach overlapped with the SNPs previously identified using EM and MCMC-based Bayesian methods, further validating the robustness of our approach. Both simulation studies and empirical data analyses showed that our method outperformed all other methods compared.

## Introduction

Genotype by environment (Q×E) interaction is present when the same cultivar (also called genotype in plant breeding) exhibits different performance across various environments. In terms of quantitative trait locus mapping, it is called QTL by environment interaction (Q×E). When data are collected from multiple environments, simply taking the mean value of the phenotypic values across multiple environments for QTL mapping will only detect the main effects of QTL. Many QTL may affect traits only through interaction with specific environments and these QTL will often be missed in the analysis where the average phenotypic values of multiple environments are used as the response variable.

Proper statistical models are required to detect QTL with Q×E interaction from data collected across multiple environments. Piepho (2000) proposed a mixed model for detecting QTL main effects from multi-environment data, while ignoring Q×E interaction QTL. Similar to composite interval mapping, his model incorporated one putative QTL at a time, along with a few cofactors (markers) selected from the whole genome. He treated the QTL effect as fixed and the environmental effects as random. Therefore, the Q×E effects were lumped into the residual as random effects. In this model, only fixed effects and variance components are considered as parameters, reducing the number of model parameters by treating the Q×E interaction variance as one of the parameters.

Boer et al. (2007) proposed a step-by-step mixed-model approach to detect QTL main effects, Q×E interaction effects and QTL responses to specific environmental covariates. In the final step, Boer et al. (2007) rewrote the model to include all QTL in a multiple-QTL framework and re-estimated their effects. Li et al. (2015) also employed a two-step approach to identify Q×E interaction in multiple environments. The first step involved using a stepwise regression within each environment to identify the significant markers, which were then used to adjust the phenotypic values. In the second step, these adjusted values from all environments were treated as the phenotypic data for QTL mapping via the inclusive composite interval mapping (ICIM). This is a way of controlling for the genetic background. In our previous studies, we proposed a multiple QTL model for Q×E interaction where all QTL effects are included in the same model (Chen et al., 2010; Zhao and Xu, 2012a, b). The method may be the optimal one because of the multiple QTL nature. However, it is not realistic to use the model when the number of markers is extremely large. To align QTL mapping for Q×E interaction with the technology of genome-wide association studies (GWAS), here we try to simplify the method via a genome scanning approach by testing one marker at a time. Such a genome scanning approach can handle virtually unlimited number of markers.

Recently, Kang *et al*. (2014) adopted a meta-analysis approach in GWAS to test Q×E interaction across multiple environments. Meta-analysis is a statistical synthesis of information from multiple independent studies to increase power and reduce false positives. It utilizes summary results from each study and thus does not require sharing individual-level information. This approach has become widely used for discovering new genetic loci associated with common diseases in human genetics. Meta-analysis in GWAS can take advantage of the new waves of even more extensive data accumulated from sequencing efforts (Evangelou and Ioannidis, 2013). Kang *et al*. (2014) treated each environment as an independent study and used the test result from each environment as the input data. These environment-specific results are then pooled to generate a final consensus test statistic.

Although the aforementioned meta-analysis accounts for the genomic background if the intermediate results are obtained from a polygenic background control model, the genomic background due to Q×E interaction has been ignored. It is well known that if the polygenic background effect is neglected, the GWAS will have an inflated residual error variance, which may reduce the statistical power and lower the resolution of GWAS (Wang et al., 2016; Wen et al., 2019; Xu, 2013; Zhou et al., 2022). We believe that the lack of the Q×E background control will cause loss of statistical power for traits primarily controlled by polygenic effects. The classical mixed model GWAS detects only additive marker effect. The genomic background effect can be controlled by fitting the kinship matrix drawn from pedigree information (if available) or inferred from genome-wide marker information (if pedigree data is unavailable). Thus, the primary goal of this study is to develop a novel linear mixed model that simultaneously detects both the main and the Q×E interaction effects of QTL, while controlling for both the main and the interaction background effects.

## Results

### Simulation study

Before analyzing the real data of rice, we present the result of simulation to show the difference among the new mixed model, the meta-analysis, the ICIM method and the fixed model (without polygenic background control). We first evaluate the null model (no marker effects are fitted to the model) for estimating three variance components, including two polygenic variances (the main effect variance and the interaction variances) and one residual variance. The model is

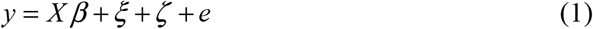

The simulation was conducted based on the actual rice genome of the RIL population so the sample size is fixed at 210 lines and the genotypes are fixed at 1619 synthetic markers (bins). Five bins were assigned as QTL whose effects are listed in **Table 1**. Variances of the polygenic main effect and polygenic Q×E interaction effect are 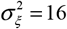 and 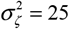, respectively, and the residual error variance is *σ* ^2^ = 9 . The means of the four populations are *μ* = {0, 2, 4, 6} . All parameters used in the simulation are given in **Table 1**. Of the five bins, only one bin (bin1300) has no Q×E interaction effect because the QTL effect is 10 across all four environments. Bin500 has no main effect because the average QTL effect over the four environments is zero.

**Table 1.** QTL effects and variance parameters used in the simulation study.

| Bins | Environ1 | Environ2 | Environ3 | Environ4 | $\sigma_{\xi}^2$ | $\sigma_{\zeta}^2$ | $\sigma^2$ | $h_{Genomics}^2$<sup>a</sup> |
| --- | --- | --- | --- | --- | --- | --- | --- | --- |
| Bin100 | 18 | 11 | 15 | 12 |  |  |  |  |
| Bin500 | -5 | 3 | 5 | -3 |  |  |  |  |
| Bin1000 | 12 | 8 | 3 | 16 | 16 | 25 | 9 | 0.775 |
| Bin1300 | 10 | 10 | 10 | 10 |  |  |  |  |
| Bin1600 | 8 | 13 | 20 | 12 |  |  |  |  |

<sup>a</sup> $h_{Genomic}^2 = (\sigma_{\xi}^2 + \sigma_{\zeta}^2) / (\sigma_{\xi}^2 + \sigma_{\zeta}^2 + \sigma^2)$ is called the genomic heritability.

**Table 2.** Variance components estimated from the new method for four traits from the 210 recombinant inbred lines of rice.

| Trait | *Mean* | | | | $\sigma_{\xi}^2$ | $\sigma_{\zeta}^2$ | $\sigma^2$ | $h_{\text{Genomics}}^2$<br>a |
| --- | --- | --- | --- | --- | --- | --- | --- | --- |
|  | 1997 | 1998 | 1999 | 2000 |  |  |  |  |
| KGW | 20.19±3.23 | 22.49±4.15 | 30.61±7.01 | 30.26±7.74 | 31.02 | 16.28 | 21.79 | 0.6846 |
| TILLER | 7.78±1.21 | 10.81±1.70 | 13.41±1.78 | 11.62±1.60 | 2.13 | 0.066 | 1.46 | 0.6001 |
| GRAIN | 107.89±22.51 | 87.40±17.34 | 93.43±21.91 | 108.95±23.38 | 1075.26 | 656.75 | 125.44 | 0.9325 |
| YIELD | 24.72±2.49 | 24.25±2.96 | 24.70±2.66 | 23.95±2.58 | 29.57 | 17.74 | 21.79 | 0.6847 |

<sup>a</sup> $h_{\text{Genomic}}^2 = (\sigma_{\xi}^2 + \sigma_{\zeta}^2) / (\sigma_{\xi}^2 + \sigma_{\zeta}^2 + \sigma^2)$ is called the genomic heritability.

The simulation experiment was replicated 100 times. Each sample was analyzed with both the new mixed model procedure and the meta-analysis approach. Furthermore, we compared the fixed model procedure and the ICIM method for the simulated data.

The average −Log_10_ (*P*) statistics across the 100 replicated samples plotted against genome locations are shown in **Figure 1**. All bins overlapping with the true effects were detected by the new mixed model method, but none of them were detected by the meta-analysis. The second peak appears at bin500, which was detected as a Q×E interaction locus with a power of 94%. On the other hand, the fourth peak at bin1300 was detected as a main effect with a power of 91%. The remaining three peaks are contributed by both the main effects and the Q×E interaction effects. Furthermore, we summarized the false positive rates across 884 bins spanning chromosomes 2, 4, 5, 7, 8, 9, and 11, where no QTLs with main or Q×E interaction effects were present. For the main effects, the false positive rate was 0.0%. In contrast, 45 bins showed false positive rates for Q×E interaction, with 33 bins exhibiting a 1% rate, 6 bins a 2% rate, 5 bins a 3% rate, and 1 bin a 4% rate. The power of ICIM method was lower than the new mixed model but higher than the meta-analysis. It should be noted that the ICIM method scans the intervals between adjacent markers rather than testing the marker positions. As a result, the peaks appeared around the target loci and produced higher false positive rates. The conclusion from this simulation study is that the new mixed model procedure outperforms the current meta-analysis and the ICIM method. The fixed model approach can detect main effects with a power of 100%, but the power to detect Q×E interaction effects is quite low.

**Figure 1.**
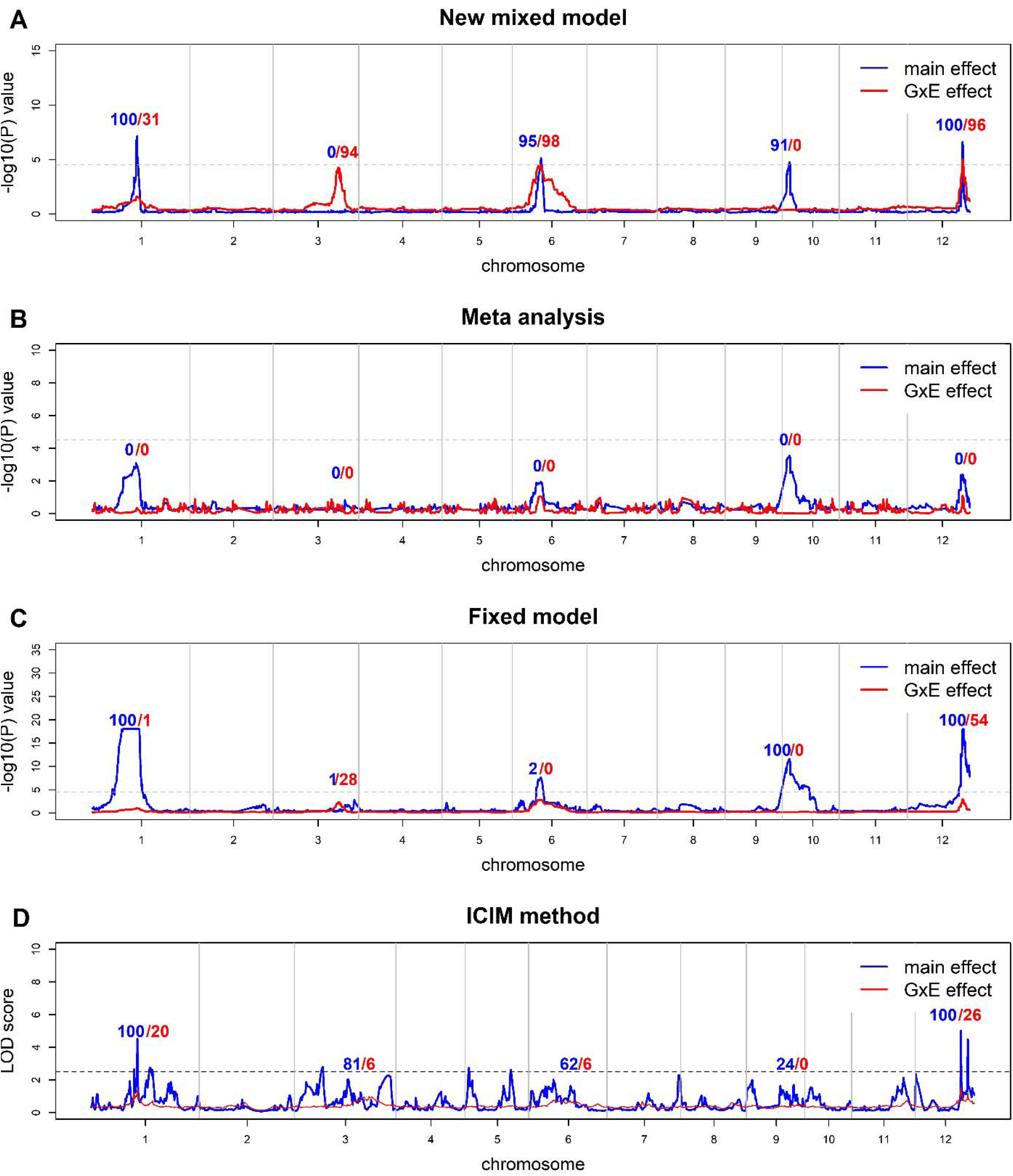
Average test statistic −Log_10_ (*P*) plotted against genome location from 100 replicated simulations. The top panel (A) is the plot for the new mixed model developed in this study. The panel (B) is the plot for the meta-analysis. The panel (C) is the plot for the fixed model (without polygenic background control). The horizontal line indicates the Bonferroni corrected threshold −Log_10_ (0.05 / 1619) = 4.51 , above which significance is declared. If the −Log_10_ (*P*) >20, the value is set to 20. The panel at the bottom (D) is the plot for the ICIM method. The horizontal line indicates threshold LOD=2.5. The blue curve represents the test for main effect and the red curve indicates the test for Q×E interaction. The numerical numbers above the peaks are the statistical powers (%) out of 100 replicates. For example, the second peak on the top panel has 0% power for the main effect and 96% power for the Q×E interaction.

### Empirical data analysis

#### The Rice data

**Table 3** shows the estimated parameters for the four traits of the rice population. The total phenotypic variance contributed by the genomic information is called the genomic heritability (de los Campos et al., 2015), which is defined as

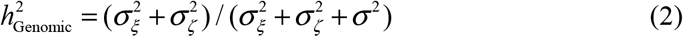

**Table 3.** Summary of all QTL regions detected by the new mixed model analysis for the four traits of rice.

| Trait | Effects | Chr | Bins | Pos(Mb) | Gene | Reference |
| --- | --- | --- | --- | --- | --- | --- |
| YIELD | Interaction | 7 | Bin975 | 0.701-0.821 |  |  |
|  | Interaction | 7 | Bin994-Bin1023 | 4.972-19.259 | *PROG1, Ghd7* | Tan et al. 2008; Xue et al. 2008 |
| GRAIN | Interaction | 6 | Bin844 | 2.325-2.415 |  |  |
|  | Interaction | 6 | Bin855-Bin859 | 3.291-4.022 | *d3* | Ishikawa et al. 2005 |
|  | Interaction | 7 | Bin995-Bin1022 | 6.027-18.805 | *PROG1, Ghd7* | Tan et al. 2008; Xue et al. 2008 |
| KGW | Main | 5 | Bin727-Bin729 | 3.973-5.376 | *GW5/qsw5* | Liu et al. 2017;Wen et al. 2018 |
|  | Interaction | 4 | Bin652-Bin654 | 29.494-29.969 |  |  |
|  | Interaction | 7 | Bin994 | 4.972-6.027 |  |  |
|  | Interaction | 7 | Bin998-Bin1023 | 6.962-19.259 | *PROG1, Ghd7* | Tan et al. 2008; Xue et al. 2008 |

All traits except KGW (the first trait) show greater contributions from the main effects than from the Q×E interaction effects. The two traits, KGW and GRAIN, have higher genomic heritability than YIELD and TILLER. The results are consistent with the conventional heritability (in ranking) except that the genomic heritability is substantially higher than the conventional heritability estimated from the replicated experiments. Issues related to the genomic heritability will be discussed again later.

In addition to the mixed model, the meta-analysis and the ICIM method, we also investigated the fixed effect model in which the polygenic background information is completely ignored. **Figure 2** shows the test statistic −Log_10_ (*P*) plotted against genome location for trait YIELD from the four methods: the new mixed model (panel A), the meta-analysis (panel B), the fixed model (panel C) and the ICIM method (panel D). The mixed model method detected one region with a significant Q×E interaction effect on chromosome 7 (panel A). This region was also detected by the meta-analysis, which detected an additional QTL on chromosome 3 with Q×E interaction effect (panel B). The fixed model detected the same region on chromosome 7 with both the main and the Q×E interaction effects (panel C). In addition to this region, the fixed model detected many regions with significant main effects. The fixed model approach does not properly control the polygenic background effect and the additional peaks may be just false positives. The ICIM method identified only one region on chromosome 1 with a significant main effect.

**Figure 2.**
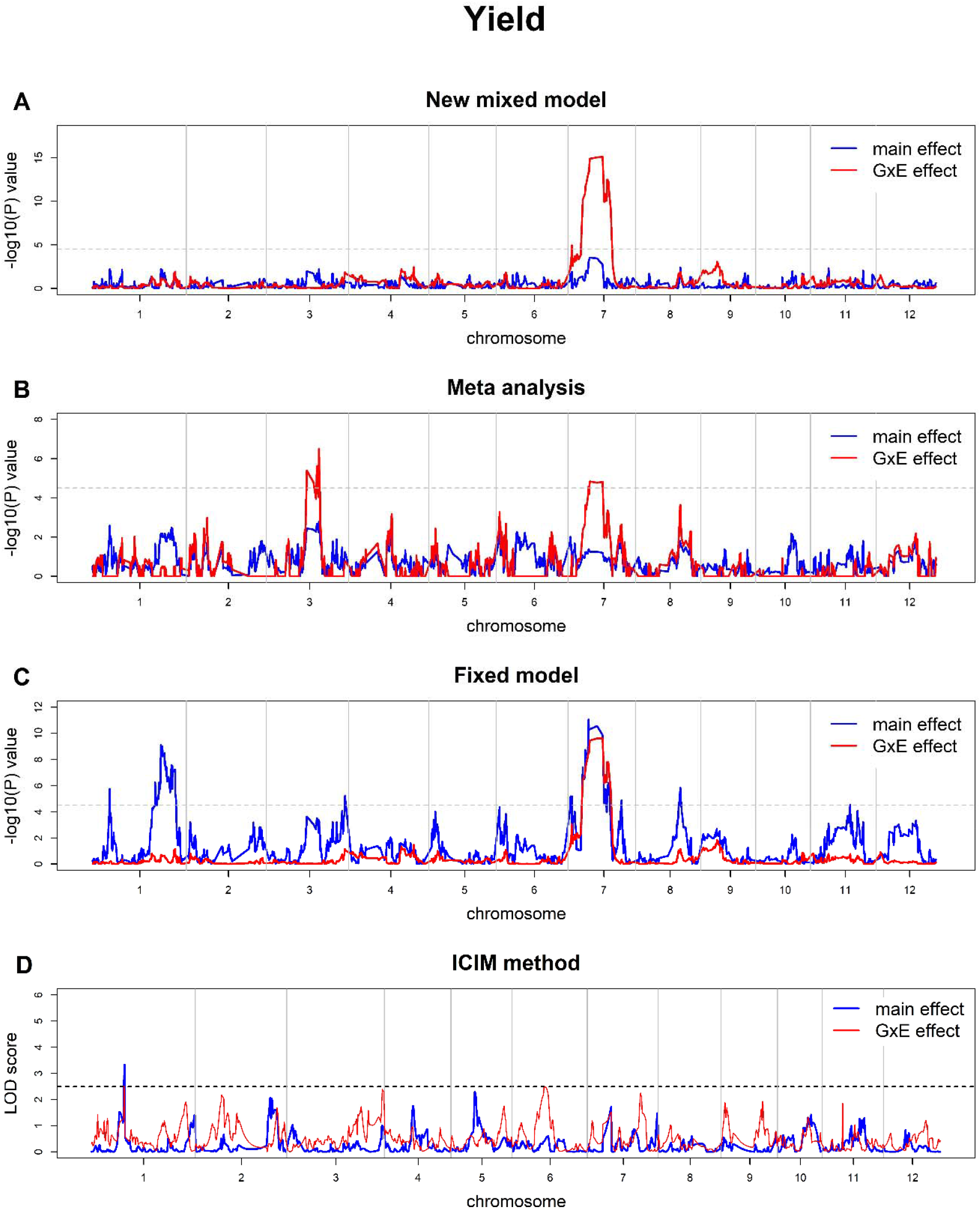
Plot of the test statistic −Log_10_ (*P*) against genome location for yield (YIELD) in the RIL population of rice. The top panel (A) is the plot from the mixed model developed in this study. The panel (B) is the plot from the meta-analysis. The panel (C) is the plot from the fixed model (without polygenic background control). The horizontal line indicates the Bonferroni corrected threshold −Log_10_ (0.05 / 1619) = 4.51 , above which significance is declared. The panel at the bottom (D) is the plot for the ICIM method. The horizontal line indicates threshold LOD=2.5. The blue curve represents the test for the main effect and the red curve indicates the test for the Q×E interaction.

Figure 3 shows the test statistic −Log_10_ (*P*) plotted against the genomic location for trait TILLER from the four methods. There are no genomic regions showing any significant effects (panel A). Although one region on chromosome 7 shows a little bump for TILLER by the new mixed model, the peak value of −Log_10_ (*P*) is 4.36, which is close to but not yet reach to the Bonferroni corrected threshold −Log_10_ (0.05 / 1619) = 4.51 . This region was identified by the ICIM method with a significant Q×E interaction effect. The meta-analysis detected nothing (panel B). The fixed model detected a main effect in the same region on chromosome 7 as the one detected by the mixed model. Many other regions are also declared as significant with the main effects by the fixed model approach. Again, these additional peaks may just be false positives. The ICIM method identified six regions with main effects and two regions with Q×E interaction effects (panel D).

**Figure 3.**
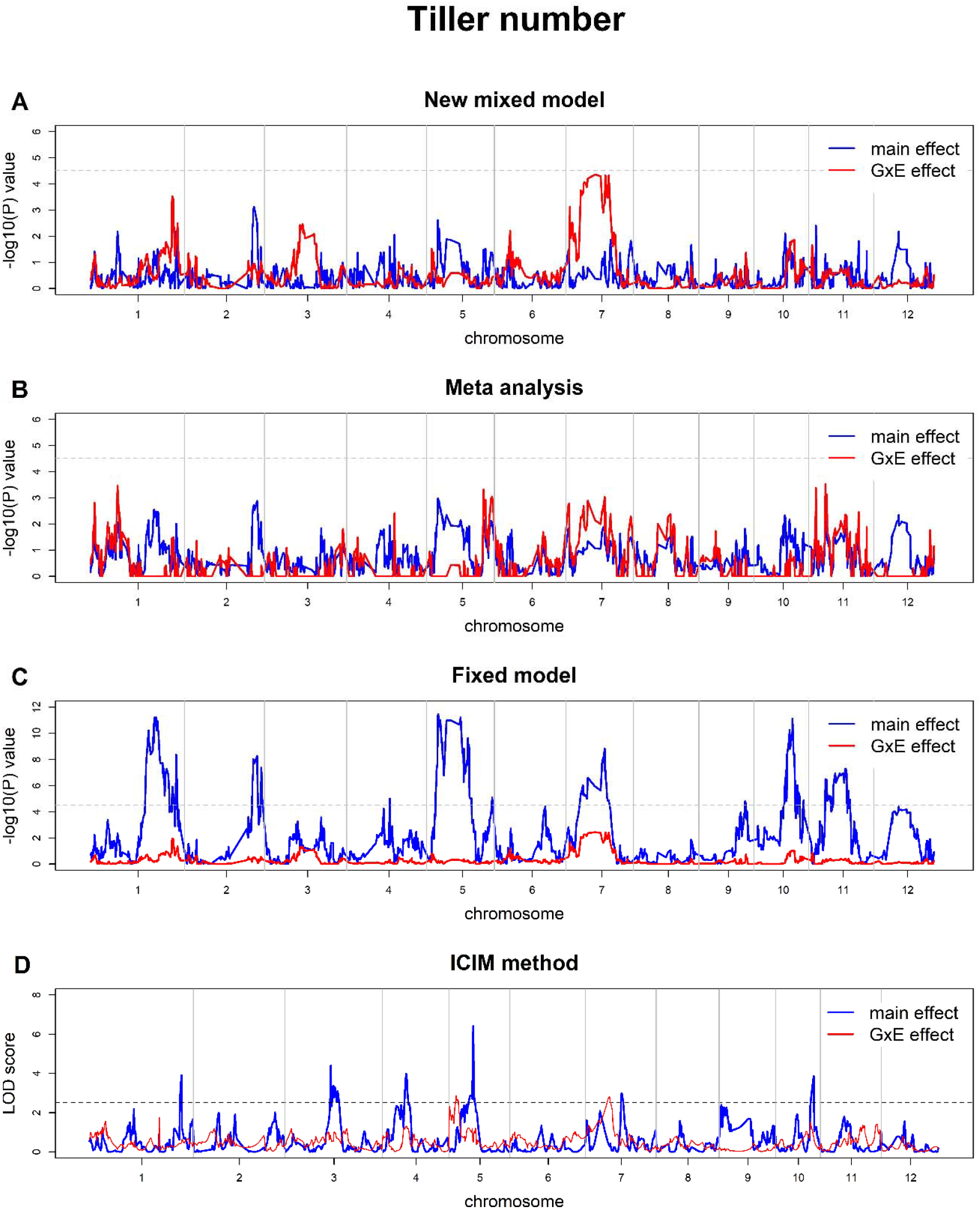
Plot of the test statistic −Log_10_ (*P*) against genome location for tiller number (TILLER) in the RIL population of rice. The top panel (A) is the plot from the mixed model developed in this study. The panel (B) is the plot from the meta-analysis. The panel (C) is the plot from the fixed model (without polygenic background control).The horizontal line indicates the Bonferroni corrected threshold −Log_10_ (0.05 / 1619) = 4.51 , above which significance is declared. The panel at the bottom (D) is the plot for the ICIM method. The horizontal line indicates threshold LOD=2.5. The blue curve represents the test for the main effect and the red curve indicates the test for the Q×E interaction.

Plots of the test statistic −Log_10_ (*P*) against the genomic location for trait KGW are shown in **Figure 4** from the four methods: the new mixed model (panel A), the meta-analysis (panel B), the fixed model (panel C) and the ICIM method (panel D). The new mixed model detected one QTL region for the main effect (chromosome 5) and three QTL regions on chromosomes 4 and 7 for the Q×E interaction effects (panel A). The meta-analysis detected two QTL regions (chromosomes 1 and 5) with Q×E effects only and these two regions are entirely different from the Q×E interaction regions detected from the mixed model (panel A). The fixed model detected many regions with main effects but none with the Q×E interaction effects. In contrast, the ICIM method detected multiple regions with main effects along with two regions on chromosome 11 exhibiting the Q×E interaction effect (panel D).

**Figure 4.**
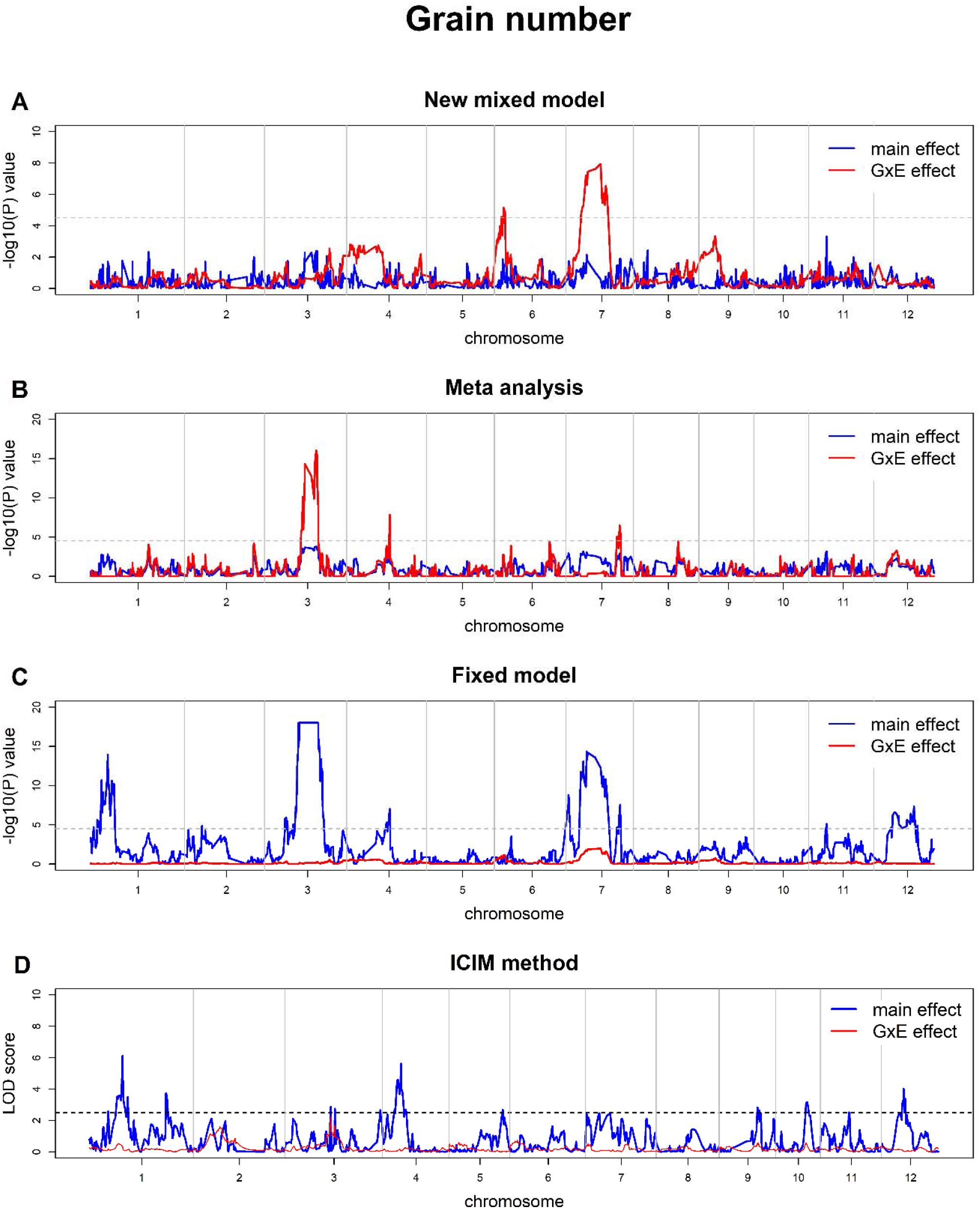
Plot of the test statistic −Log_10_ (*P*) against genome location for grain number (GRAIN) in the RIL population of rice. The top panel (A) is the plot from the mixed model developed in this study. The panel (B) is the plot from the meta-analysis. The panel (C) is the plot from the fixed model (without polygenic background control). If the −Log_10_ (*P*) >20, the value is set to 20. The horizontal line indicates the Bonferroni corrected threshold −Log_10_ (0.05 / 1619) = 4.51 , above which significance nis declared. The panel at the bottom (D) is the plot for the ICIM method. The horizontal line indicates threshold LOD=2.5. The blue curve represents the test for the main effect and the red curve indicates the test for the Q×E interaction.

Results for trait GRAIN are shown in **Figure 5**. The new mixed model method detected two QTL regions (chromosomes 6 and 7) with Q×E interaction effects (panel A). The meta-analysis identified three Q×E interaction regions (chromosomes 3, 4 and 7), which are different from the regions detected by the new mixed model. In contrast, the fixed model detected many regions with main effects only. A similar pattern was observed with the ICIM method.

**Figure 5.**
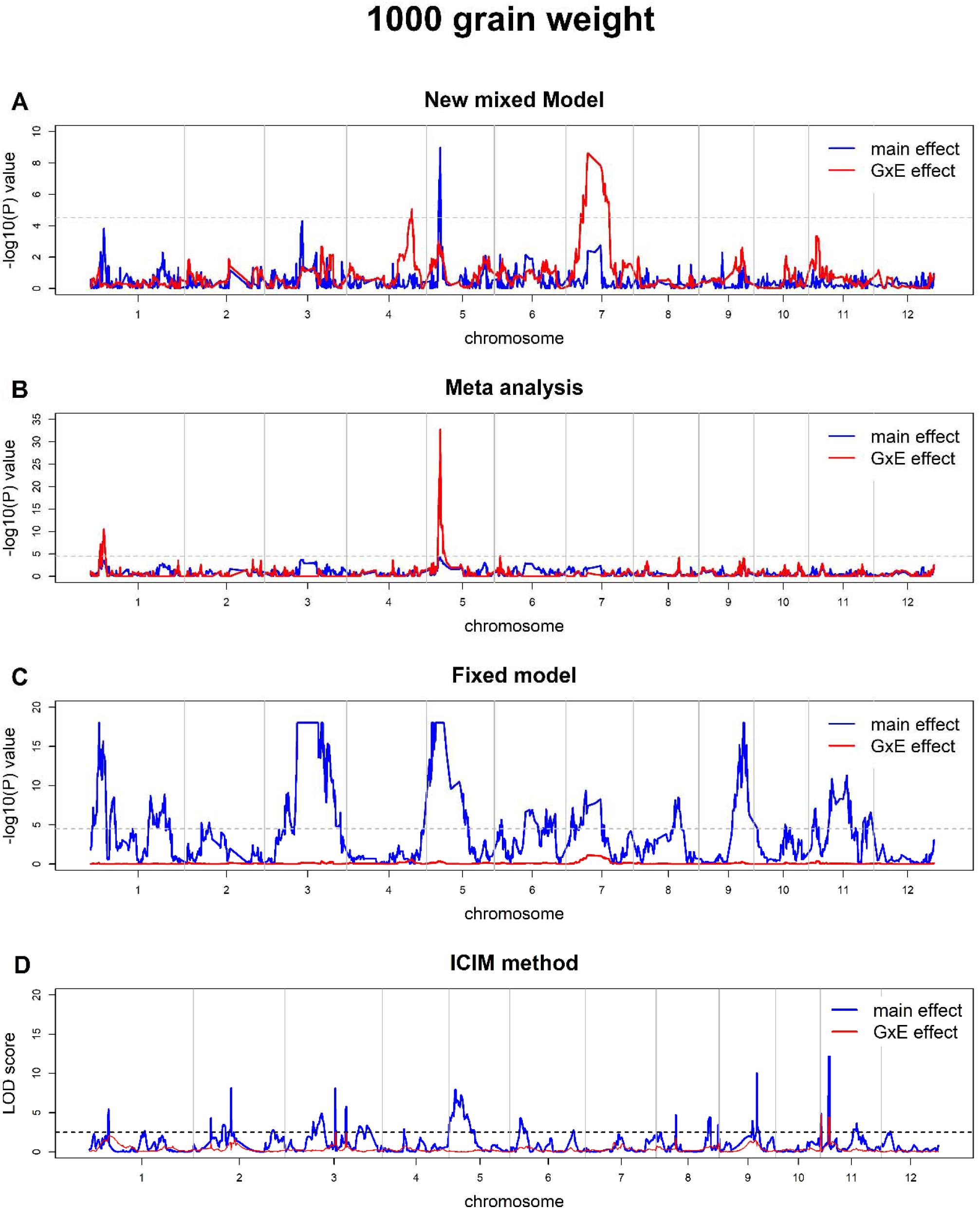
Plot of the test statistic −Log_10_ (*P*) against genome location for 1000 grain weight (KGW) in the RIL population of rice. The top panel (A) is the plot from the mixed model developed in this study. The panel (B) is the plot from the meta-analysis. The panel (C) is the plot from the fixed model (without polygenic background control).The horizontal line indicates the Bonferroni corrected threshold −Log_10_ (0.05 / 1619) = 4.51 , above which significance is declared. If the −Log_10_ (*P*) >18, the value is set to 18. The panel at the bottom (D) is the plot for the ICIM method. The horizontal line indicates threshold LOD=2.5. The blue curve represents the test for the main effect and the red curve indicates the test for the Q×E interaction.

Additionally, we provide the estimates of main and Q×E interaction effects for four traits of rice in **Supplementary File S1**. It should be noted that we measured the variance of environment specific effects of the locus under investigation as in our previous studies (Chen et al., 2010; Zhao and Xu, 2012a, b). From the simulation study and real data analyses, we conclude that the fixed model (without properly controlling the polygenic background) has no power to detect Q×E interaction effects. **Table 4** summarizes all QTL regions detected from the new mixed model for the four traits. Genes with known functions in previous studies that are located on the genomic regions detected here are also shown in **Table 4**. Of the four traits, only KGW has a main effect QTL located on chromosome 5 and it was detected only by the mixed model. This genomic region contains a known QTL GW5/qsw5 (Liu et al., 2017). The two methods (mixed model and meta-analysis) detected a total of six Q×E interaction QTL for YIELD, another six Q×E interaction QTL for GRAIN and seven Q×E interaction QTL for KGW (**Table 4**). It appears that the QTL on chromosome 7 may have pleiotropic effects (one QTL controlling multiple traits). The genomic regions harbor two important genes (*PROG1* and *Ghd7*). The genomic regions also contain a known gene *GS3* for YIELD and GRAIN. In addition, there are other functional genes located in these regions, e.g., *d3* and *GIF1* for GRAIN and *Gn1a* for KGW. All candidate genes are listed in **Supplementary File S2**. These results demonstrated that our new method can be identified effectively QTL with the main effect, Q×E interaction effect or both.

#### The barley data

Figure 6. displays the results of QTL mapping for lodging resistance (a quantitative trait) in barley. The new mixed model identified one QTL region on chromosome 3 with both the main and Q×E interaction effects, along with additional five QTL regions exhibiting only Q×E interaction effects (chromosome 2, 3, 4, 6 and 7) (panel A). The meta-analysis detected only three Q×E interaction regions (chromosomes 2 and 4), which overlapped with the regions identified by the new mixed model. The fixed model also detected the regions with the Q×E interaction effect on chromosome 2. In comparison, the ICIM method detected three regions (chromosomes 2 and 3) with the Q×E interaction effect and four regions (chromosomes 3, 4 and 7) with main effects.

**Figure 6.**
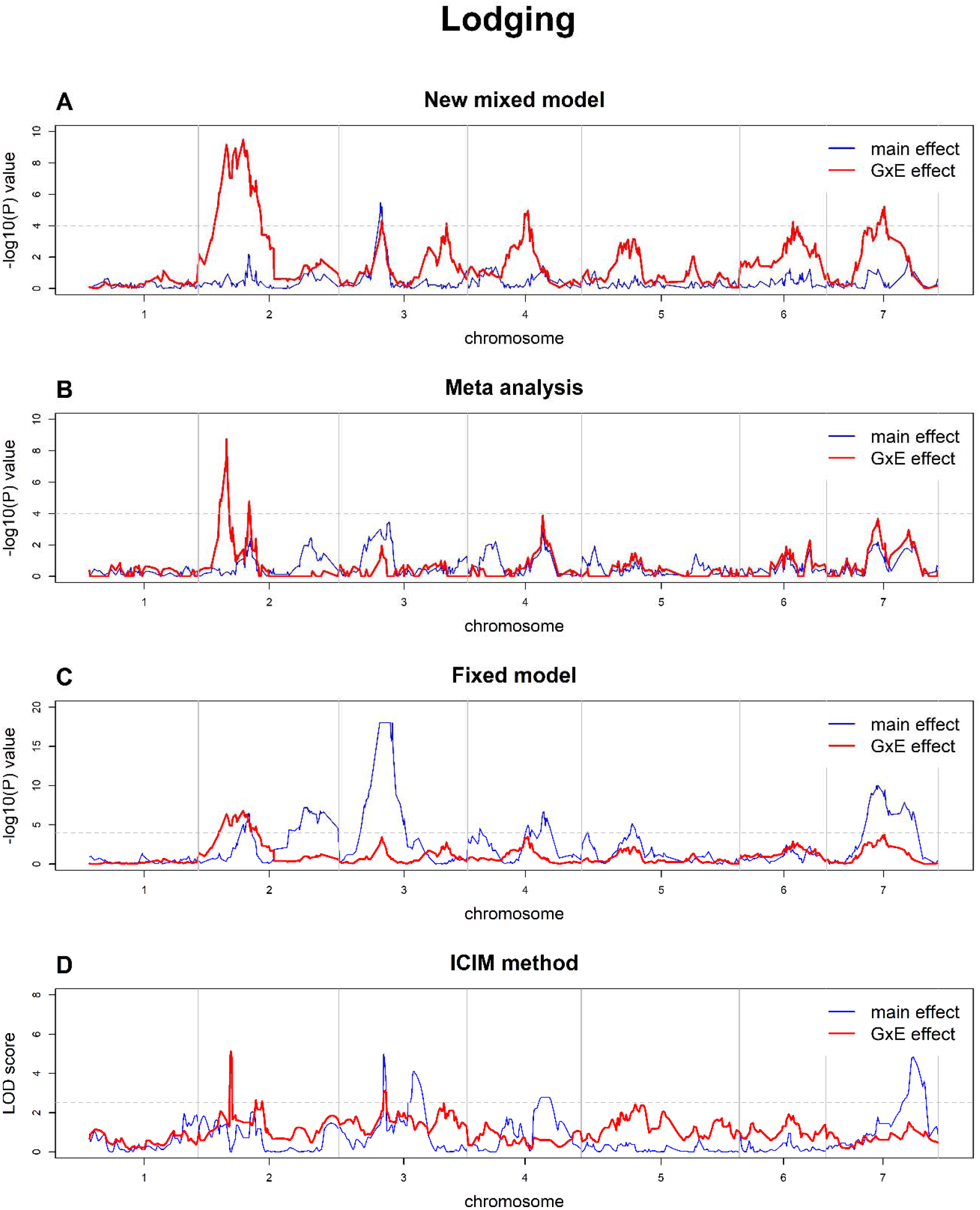
Plot of the test statistic −Log_10_ (*P*) against genome location for lodging in the doubled haploid of barley. The top panel (A) is the plot from the new mixed model developed in this study. The panel (B) is the plot from the meta-analysis. The panel (C) is the plot from the fixed model (without polygenic background control). The horizontal line indicates the Bonferroni corrected threshold −Log_10_ (0.05 / 1619) = 4.51 , above which significance is declared. If the −Log_10_ (*P*) >18, the value is set to 18. The panel at the bottom (D) is the plot from ICIM method. The horizontal line indicates the LOD threshold LOD=2.5.The blue curve represents the test for the main effect and the red curve indicates the test for the Q×E interaction.

## Discussion

The unique feature of the linear mixed model procedure for Q×E interaction is the way to control the polygenic background information. We defined two separate polygenic backgrounds captured by two different kinship matrices, the kinship matrix for the main effects 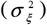 and the kinship matrix for the Q×E interactions 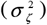. With such a polygenic background control, many false positives can be prevented with the linear mixed model methodology. In contrast, the ICIM method controls the background effects by using selected markers across the genome as covariates.

Although the polygenic background here played the same role as cofactors in ICIM method, the kinship matrix was constructed using the genome-wide marker information rather than a limited set of markers. To reduce the number of variance components in the mixed model, Zhou et al. (2022) combined both the main and Q×E interaction polygenic effects to create a compressed polygenic background. In our rice data analysis, we observed many significant regions for the main effects with the fixed model approach where the polygenic background is not properly controlled. We also noticed that the fixed model approach detected only one region with significant Q×E on chromosome 7 for YIELD, while the significant Q×E regions on additional chromosomes for other traits detected by the other methods for other traits are completely missed by the fixed model. In the barley data analysis, all six regions exhibiting Q×E interaction effects identified by the new mixed model overlapped with the SNPs identified using the EM and MCMC-based Bayesian methods in our previous study (Zhao and Xu, 2012a).

In this study, the genomic relationship matrix (GRM) for each environment was constructed using the following equation 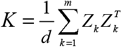, where *Z*_*k*_ is the genotype vector at the *k*th locus, *m* is the total number of SNPs and *d* is the summation of diagonal elements of *K*. This GRM is similar to those proposed by VanRaden (2008), which makes the polygenic background control in this study comparable to the genomic best linear unbiased prediction where the additive kinship matrix is used.

The methods for constructing the GRM can be used either the genome-wide SNPs (VanRaden, 2008; Xu, 2013) or ROHs (Zhao et al., 2023). In our previous studies, we observed these GRM construction methods influenced prediction accuracies in genomic selection (Li et al., 2025a). Additionally, kernel-based kinship matrices, such as the correlation matrix, have been used in genomic selection (Cuevas et al., 2016; Li et al., 2025b). How the different GRM and kernel-based kinship matrices perform for the Q×E interaction study is certainly an interesting future topic.

The meta-analysis adopted here utilizes intermediate results from environment specific studies, *i*.*e*., the estimated QTL effect and the estimation error for each locus. These intermediate results are then treated as input data for a second model involved in the meta-analysis. Such a meta-analysis allows us to separate the Q×E interaction effect from the main effect. If one is not interested in Q×E interaction but only interested in combining multiple environmental data to improve statistical power, a much simpler meta-analysis is available. For example, the Fisher’s method (Fisher, 1932) only uses the final results of separate studies, the *p*-values, for meta-analysis.

Let *p*_*t*_ be the *p*-value from the *t*th environment for the locus of interest. Fisher’s consensus test statistic is

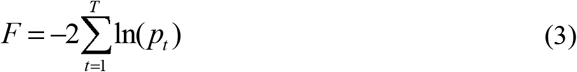

Under the null model, *i*.*e*., the QTL effect is absent in all environments, the consensus test statistic *F* follows a Chi-square distribution with 2*T* degrees of freedom. We also used this test to combine test results from all four environments and the results are given in **Supplementary File S3**.

Before scanning the entire genome for Q×E interaction effects, we estimated the genomic heritability for each of the four traits. Genomic heritability is defined as the proportion of phenotypic variance contributed by all genome-wide markers. This parameter does not provide much useful information other than allows the investigators to have a general idea of how the program behaves. If the estimated genomic heritability is zero, the program or the data may be problematic. On the other hand, if the genomic heritability is one, it may reflect an overfitting (very often when the number of markers is huge). One should not take the genomic heritability seriously because a genomic heritability of unity does not mean the trait being 100% controlled by genes. More discussion on genomic heritability can be found in de los Campos et al. (2015). We actually estimated the heritability for each of the four traits from the replicated data (four environments) using the conventional analysis of variances. The results are shown in **Supplementary File S4** and these estimates are indeed much lower than the genomic heritability. Although overfitting observed, it does not affect tests for the main effects and the Q×E interaction effects. This is because the tests rely on general error consisting of all random effects and the residual errors. Overfitting only affects the relative distribution (or partitioning) of different variance components and does not affect the total variance (Xu et al., 2016).

To the best of our knowledge, this is the first study to identify Q×E interaction for these four traits in this RIL population. We did not found any main effect QTL and any Q×E interaction effect QTL for TILLER. As shown in **Table 4**, several important functional genes have been isolated and cloned on the genomic regions detected in this study. *qSW5/GW5* is a major gene for grain width and grain weight in rice, and a major determinant of grain yield in cereal crops (Liu et al., 2017). *D3* can regulate the number of tillers (Ishikawa et al., 2005), which can indirectly control the grain number (GRAIN). It should be noted that one genomic region on chromosome 7 has been detected by the new mixed model for three of the four traits (YIELD, GRAIN and KGW). This region harbors two important genes, *PROG1* and *Ghd7. PROG1* has been documented as being associated with erect growth, greater grain number and higher grain yield in cultivated rice (Jin et al., 2008). It may control tiller angle and tiller number in rice (Tan et al., 2008). *Ghd7* may have large pleiotropic effects on an array of traits, including grain number, heading date and plant height (Xue et al., 2009). Such pleiotropic effects may provide an explanation for QTL hot spots that have been observed in many studies.

## Material and Methods Mixed model methodology

Let *y*_*t*_ be an *n* ×1 vector of phenotypic values of a trait collected from the *t*-th environment for *t* = 1,L ,*T* where *T* is the number of environments and *n* is the number of plants (sample size). Let 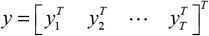 be an (*nT* ) × 1 vector holding the phenotypic values from all plants in all environments. The Q×E model is

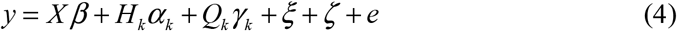

where *X* is an (*nT* ) × *T* design matrix capturing the environmental effects, *β* is an *T* ×1 vector of the *T* environment effects, *H*_*k*_ is an (*nT* ) ×1 vector of genotype indicator variable for marker *k, α* _*k*_ is the main effect (fixed effect) of the locus under investigation, *Q*_*k*_ is an (*nT* ) × *T* design matrix for Q×E interaction effects, *γ*_*k*_ is a *T* × 1 vector of environment specific effects (fixed effects) for the marker of interest, *ξ* is an (*nT* ) ×1 vector of polygenic main effects (random), *ζ* is an (*nT* ) × 1 vector of polygenic Q×E interaction effects (random) and *e* is an (*nT* ) × 1 vector of residual errors. Let *Z* be an *n*×*m* matrix holding the numerically coded genotype indicator variables where *m* is the total number of markers and n is the sample size. Matrices *H*_*k*_ and *Q*_*k*_ are some rearrangements of the *k*th column of *Z* if marker *k* is the one under consideration. The model is too complicated to understand. We provide a small example to demonstrate the elements of the model in detail (**Supplementary File S5**).

The restricted maximum likelihood estimation (REML) was employed to estimate the variance components and to obtain the best linear unbiased estimates of the fixed effects. Given the expectation of *y* and its variance, we constructed the restricted likelihood function with the assumption that the trait values are *y ∼N(μ,V)* distributed, where

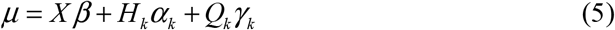

and

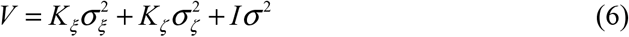

Let us define *U* = [ *X* || *H* _*k*_ || *Q*_*k*_ ] as the horizontal concatenation of the three matrices.

The restricted log likelihood function is

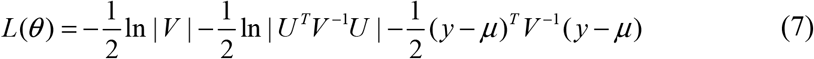

The MIXED procedure in SAS (SAS Institute, 2009) was used to estimate the parameters. Once the three variance components are estimated, 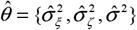, which are used to build matrix 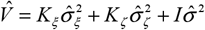, and the fixed effects are estimated via

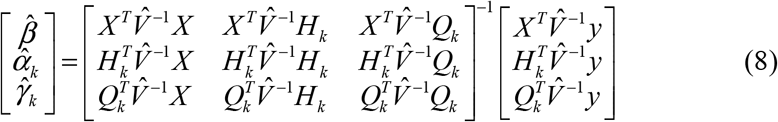

The variance matrix of the estimated fixed effects is

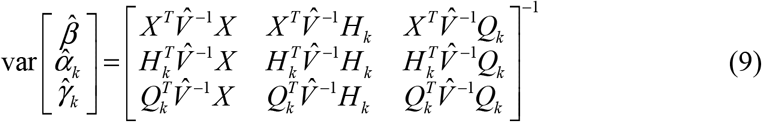

which is

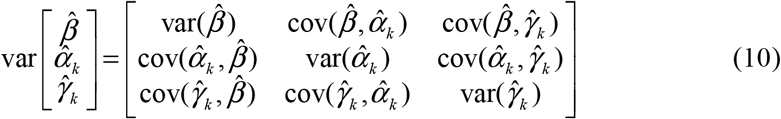

The main effect is tested based on the following Wald test

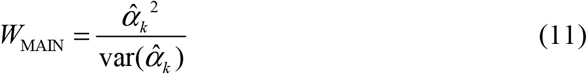

which (under the null model) follows a Chi-square distribution with one degree of freedom. The Q×E interaction effects are tested based on

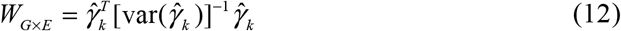

Under the null model, this Wald test statistic follows a Chi-square distribution with *T* −1 degrees of freedom. From the above Wald tests and their degrees of freedom, corresponding *p*-values are calculated, which are compared to the Bonferroni corrected threshold to determine the significance of each main effect and each Q×E interaction effect.

### Meta-analysis for Q×E interaction

Meta-analysis for Q×E requires results of GWAS from separate environments (environment specific analyses) or independent studies. For each locus, we only need the *p*-value or the estimated effect and the error of the estimate. Results of any methods of GWAS can be used in meta-analysis. However, we used the most advanced method of GWAS by controlling the polygenic background noise (Xu, 2013). When the polygenic background only contains the additive effects, the method is equivalent to the genome-wide efficient mixed model analysis for association studies (GEMMA) (Zhou and Stephens, 2012). Methods of estimating marker effects for the independent studies are not the main concerns of this study, but for the paper to be self-contained, we briefly introduce the model here. Let *y* be an *n*×1 vector for the phenotypic values of a quantitative trait from one environment. The linear mixed model is

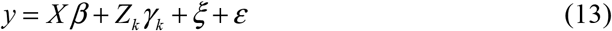

where *X* and β are the design matrix and effects for some systematic errors (treated as fixed effects); *Z* _*k*_ is a genotype indicator variable for marker *k* and *γ* _*k*_ is the effect for this marker (treated as fixed effect), where *k* =1, …, *m* and *m* is the number of markers; *ξ* is a polygenic effect following a 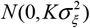 distribution, where *K* is an *n*×*n* kinship matrix calculated from genome-wide marker information and 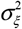 is a polygenic variance; ε is a vector of residual errors following a *N* (0, *Iσ* ^2^ ) distribution, where *σ* ^2^ is the residual error variance.

Let us denote the estimated effect and its estimation error variance for marker *k* by 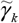 and 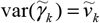, where *k* =1, …, *m* for all markers in the genome. For *T* separate studies, we will have *T* such estimates and error variances. We now focus our analysis on one marker since the genome-wide marker analysis simply loops the single-marker analysis over all markers. We now drop subscript *k* and replace it by *t* for the same marker but over *t* = 1, … , *T* environments. In other words, 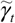 and 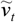 are the estimated effect and its error variance for the current marker of interest from the *t*th environment.

After genome-wide detection over all environments, we have 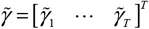 and 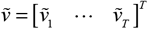 which are treated as input data to build another linear mixed model

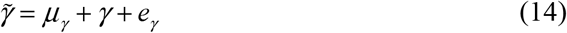

where *μ*_*γ*_ is a mean value representing the main effect for the marker of interest, γ Is a vector of true effects following an assumed 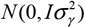 distribution, and *e*_*γ*_ is an *T*×1 vector of errors with a known 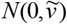 distribution, where 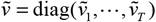.

The parameters are *μ*_*γ*_ and 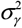 . The log likelihood function is

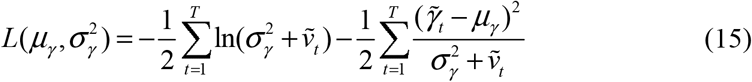

Numerical solutions can be obtained via the Newton-Raphson method. Let 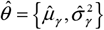 be the MLE of the parameters. We now have two hypotheses to test, one for the main effect *H*_0_ : *μ*_*γ*_ = 0 and one for the Q×E interaction 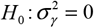. Rejection of either hypothesis means a significant association. If we are not interested in differentiation of whether the association is due to the main effect or due to the interaction effect, we can combine the two tests together to develop a composite test 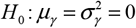. Let the likelihood value under the null model be

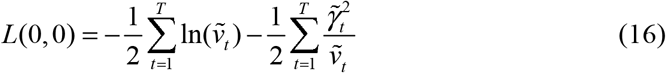

The likelihood ratio test statistic is

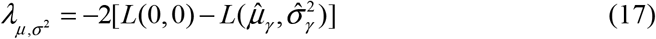

Under the null model, this test statistic follows approximately a mixture of Chi-square one and Chi-square two distributions with an equal weight (Zaman et al., 2006).

Therefore, the *p*-value is calculated using

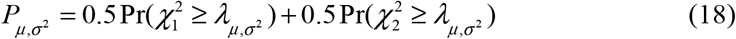

Where 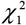 and 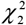 are variables of Chi-square one and Chi-square two, respectively. The approximation can be very good if *T* is relatively large.

We now introduce separate tests for the main effect (mean) and the interaction effect (variance). For the main effect hypothesis, *H*_0_ : *μ*_*γ*_ = 0 , the Wald test is

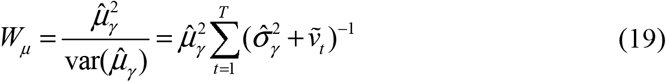

Under the null hypothesis, the above Wald test follows approximately a Chi-square distribution with one degree of freedom. For the Q×E interaction hypothesis, 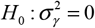, the likelihood ratio test is appropriate, which is defined as

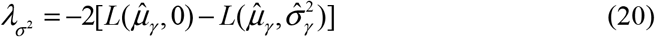

where 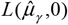 is the likelihood value evaluated at 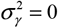 . This test statistic follows approximately a mixture distribution of 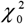 and 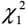 with an equal weight. Therefore, the *p*-value is calculated using

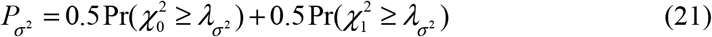

Again, the subscript (*σ* ^2^ ) in the test statistic and the *p*-value means that this test is for 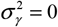. The Q×E interaction effect represented by the variance 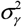.

### Experimental data

#### The rice data

The RIL population of rice was initially developed by Hua *et al*.(2003; 2002) and Xing et al. (2002) and re-genotyped by Yu et al.(2011). The population consists of 210 RILs derived from the cross of two elite inbred varieties, Zhenshan 97 and Minghui 63. The hybrid of the two varieties, Shanyou 63, was undergoing nine generations of selfing via single seed descent to generate the 210 RILs. The phenotypic values were field evaluated in 1997, 1998 and 1999 in two locations (four environments) at the Experimental Station of Huazhong Agricultural University, Wuhan, China. We analyzed four agronomic traits: yield per plant (YD), 1000 grain weight (KGW), grain number per plant (GRAIN) and tiller number per plant (TILLER). Markers are represented by 1619 bins inferred from 270,820 single nucleotide polymorphisms (SNPs) across 12 chromosomes of the rice genome. All SNPs within a bin have exactly the same segregation pattern (perfect linkage disequilibrium or LD) and thus one SNP from a bin is sufficient to represent the entire bin. Bin genotypes of the 210 RILs were coded as 1 for the Zhenshan 97 genotype and 0 for the Minghui 63 genotype.

Gene annotation: We exerted gene information from http://rice.plantbiology.msu.edu/pub/data/Eukaryotic_Projects/o_sativa/annotation_dbs/pseudomolecules/version_7.0/all.dir/all.gff3 and https://funricegenes.github.io/geneInfo.table.txt. According to the genic regions, we annotated the genomic regions that are associated with the traits of interest.

#### The barley data

To access the broad applicability of the new method, we also applied it to analyze the doubled haploid (DH) population of barley, as published by Hayes et al. (1993). The genotype and phenotype data were respectively download from the following links: http://www.genenetwork.org/genotypes/S×M.geno and http://wheat.pw.usda.gov/ggpages/S×M/phenotypes.html. This dataset consists of 150 DH lines derived from the cross of two spring barley varieties (Steptoe and Morex). The total number of markers is 495 distributed along seven chromosomes of the barley genome. After removing two markers with problematic records, 493 SNPs remained for further analysis. The genotype of each marker was coded as 1 for the Steptoe allele and -1 for the Morex allele. The database contains eight quantitative traits, with lodging resistance selected as the trait of interest for this analysis. The trait was evaluated across six environments.

#### Software implementation

All data analyses were conducted using the restricted maximum-likelihood (REML) method (Patterson and Thompson, 1971) implemented via the MIXED and GLIMMIX procedures in SAS (SAS Institute, 2009). PROC IML (SAS Institute, 2009) was used to calculate the kinship matrix. The three variance components were estimated using PROC MIXED. Genome scans were implemented using PROC GLIMMIX. The SAS codes are provided in **Supplementary File S6** for PROC MIXED and **Supplementary File S7** for PROC GLIMMIX. Furthermore, the R script based on the ASReml-R (Butler et al., 2023) was compiled into the TBtools II plugin module, provided in **Supplementary File S8**. This module can be easily integrated into TBtools II (Chen et al., 2023) for straightforward implementation. The kinship matrices of polygenic main effects and interaction effects are given in **Supplementary Files S9 and S10**, respectively. The phenotypes of the four quantitative traits across six environments are given in **Supplementary File S11**. The SNP genotypes are stored in **Supplementary File S12**.

## Data availability

All data used in the study can be downloaded as supplementary material from the journal website.

## CRediT authorship contribution statement

**Fuping Zhao:** Writing-Original draft, Conceptualization, Validation, Investigation, Data curation. proposed the methodology, participated in data analysis and drafted the manuscript. **Lixian Wang:** Investigation. **Shizhong Xu:** Writing-Review & Editing, Resources, conceived of the study, and helped to draft the manuscript.

## Acknowledgements

We thanks Dr Chen Chengjie for his assistance in building the TBtool II plugin module. This work was supported by the National Natural Science Foundations of China (No. 32172702), the State Key Laboratory of Animal Biotech Breeding (XQSWYZQZ-KFYX-4), the National Key Research and Development Program of China (2024YFF1000100, 2021YFD1301102), Zaozhuang Elite Industrial Innovation Program and Agricultural Science and Agricultural Science and Technology Innovation Program (No.ASTIP-IAS-TS-6) to F.Z. The project was also supported by the United States National Science Foundation (NSF) Collaborative Research Grant (DBI-1458515) to S.X.

## Supplementary Materials

**File S1:** The estimates of main and QxE interaction effects for four traits of rice

**File S2:** All candidate genes locate in the QTL regions identified by the new mixed model proposed by us.

**File S3:** Plot of the test statistic −Log_10_ (*P*) against genome location for YIELD, TILLER, KGW and GRAIN in the RIL population of rice by Fisher’s method.

F**ile S4:** Variance components estimated for four traits from the 210 recombinant inbred lines of rice

**File S5:** A example to demonstrate the new mixed model

**File S6:** SAS PROC MIXED for Polygenic Variance Components Analysis

**File S7:** SAS PROC GLIMMIX for mapping main and Q×E interaction effects

**File S8:** TBtool-II plugin module of Q×E effect detection

**File S9:** The kinship matrix of polygenic main effect

**File S10:** The kinship matrix of polygenic Q×E interaction effect

**File S11:** The dataset contains the phenotypic values of four traits.

**File S12:** The dataset contains the genotypes.

## References

Boer, M.P., Deanne, W., Lizhi, F., Podlich, D.W., Lang, L., Mark, C., Eeuwijk, F.A. Van,, 2007. A mixed-model quantitative trait loci (QTL) analysis for multiple-environment trial data using environmental covariables for QTL-by-environment interactions, with an example in maize. Genetics 177(3), 1801–1813. 10.1534/genetics.107.071068.

Butler, D.G., Cullis, B.R., Gilmour, A.R., Gogel, B.J., Thompson, R., 2023. ASReml-R reference manual version 4.2. VSN International Ltd, https://asreml.kb.vsni.co.uk.

Chen, C., Wu, Y., Li, J., Wang, X., Zeng, Z., Xu, J., Liu, Y., Feng, J., Chen, H., He, Y., Xia, R., 2023. TBtools-II: A “one for all, all for one” bioinformatics platform for biological big-data mining. Mol Plant 16(11), 1733–1742. 10.1016/j.molp.2023.09.010.

Chen, X., Zhao, F., Xu, S., 2010. Mapping environment-specific quantitative trait loci. Genetics 186(3), 1053. 10.1534/genetics.110.120311.

Cuevas, J., Crossa, J., Soberanis, V., Pérez-Elizalde, S., Pérez-Rodríguez, P., Campos, G.D.L., Montesinos-lopez, O.A., Burgueño, J., 2016. Genomic prediction of genotype× environment interaction kernel regression models. Plant Genome 9(3), 1–20. 10.3835/plantgenome2016.03.0024.

de los Campos, G., Sorensen, D., Gianola, D., 2015. Genomic heritability: what is it? PLoS Genet 11(5), e1005048. 10.1371/journal.pgen.1005048.

Evangelou, E., Ioannidis, J.P.A., 2013. Meta-analysis methods for genome-wide association studies and beyond. Nat Rev Genet 14(6), 379–389. 10.1038/nrg3472.

Fisher, R.A., 1932. Statistical methods for research workers. 4th edition. London: Oliver and Boyd.

Hayes, P.M., Liu, B.H., Knapp, S.J., Chen, F., Jones, B., Blake, T., Franckowiak, J., Rasmusson, D., Sorrells, M., Ullrich, S.E., Wesenberg, D., Kleinhofs, A., 1993. Quantitative trait locus effects and environmental interaction in a sample of North American barley germ plasm. Theor Appl Genet 87(3), 392–401. 10.1007/BF01184929.

Hua, J., Xing, Y., Wu, W., Xu, C., Sun, X., Yu, S., Zhang, Q., 2003. Single-locus heterotic effects and dominance by dominance interactions can adequately explain the genetic basis of heterosis in an elite rice hybrid. Proc Natl Acad Sci USA 100(5), 2574–2579. 10.1073/pnas.0437907100.

Hua, J., Xing, Y., Xu, C., Sun, X., Yu, S., Zhang, Q., 2002. Genetic dissection of an elite rice hybrid revealed that heterozygotes are not always advantageous for performance. Genetics 162(4), 1885–1895. 10.1093/genetics/162.4.1885.

Ishikawa, S., Maekawa, M., Arite, T., Onishi, K., Takamure, I., Kyozuka, J., 2005. Suppression of tiller bud activity in tillering dwarf mutants of rice. Plant Cell Physiol 46(1), 79–86. 10.1093/pcp/pci022.

Jin, J., Huang, W., Jp Yang, J., Shi, M., Zhu, M., Luo, D., Lin, H., 2008. Genetic control of rice plant architecture under domestication. Nat Genet 40(11), 1365–1369. 10.1038/ng.247.

Kang, E.Y., Han, B., Furlotte, N., Joo, J.W.J., Shih, D., Davis, R.C., Lusis, A.J., Eskin, E., 2014. Meta-analysis identifies gene-by-environment interactions as demonstrated in a study of 4,965 mice. PLoS Genet 10(1), e1004022. 10.1371/journal.pgen.1004022.

Li, M., Shi, L., MachHugh, D.E., Wang, X., Tian, J., Wang, L., Deng, Y., Wang, L., Zhao, F., 2025a. Genomic prediction based on unbiased estimation of the genomic relationship matrix in pigs. Animal 19(Online), 101402. 10.1016/j.animal.2024.101402.

Li, M., Tall, T., MachHugh, D.E., Chen, L., Garrick, D., Wang, L., Zhao, F., 2025b. KPRR: A novel machine learning approach for effectively capturing nonadditive effects in genomic prediction. Brief Bioinform 26, bbae683. 10.1093/bib/bbae683.

Li, S., Wang, J., Zhang, L., 2015. Inclusive composite interval mapping of qtl by environment interactions in biparental populations. PLoS One 10(7), e0132414. 10.1371/journal.pone.0132414.

Liu, J., Chen, J., Zheng, X., Wu, F., Lin, Q., Heng, Y., Tian, P., Cheng, Z., Yu, X., Zhou, K., 2017. GW5 acts in the brassinosteroid signalling pathway to regulate grain width and weight in rice. Nat Plants 3, 17043. 10.1038/nplants.2017.43.

Patterson, H.D., Thompson, R., 1971. Recovery of inter-block information when block sizes are unequal. Biometrika 58(3), 545–554. 10.1093/biomet/58.3.545.

Piepho, H.P., 2000. A mixed-model approach to mapping quantitative trait loci in barley on the basis of multiple environment data. Genetics 156(4), 2043. 10.1093/genetics/156.4.2043.

SAS Institute, 2009. SAS 9.4 product documentation. SAS Institute, Cary, NC, USA.

Tan, L., Li, X., Liu, F., Sun, X., Li, C., Zhu, Z., Fu, Y., Cai, H., Wang, X., Xie, D., 2008. Control of a key transition from prostrate to erect growth in rice domestication. Nat Genet 40(11), 1360. 10.1038/ng.197.

VanRaden, P.M., 2008. Efficient methods to compute genomic predictions. J Dairy Sci 91(11), 4414–4423. 10.3168/jds.2007-0980.

Wang, S.B., Wen, Y.J., Ren, W.L., Ni, Y.L., Zhang, J., Feng, J.Y., Zhang, Y.M., 2016. Mapping small-effect and linked quantitative trait loci for complex traits in backcross or DH populations via a multi-locus GWAS methodology. Sci Rep 6, 29951. 10.1038/srep29951.

Wen, Y.J., Zhang, Y.W., Zhang, J., Feng, J.Y., Dunwell, J.M., Zhang, Y.M., 2019. An efficient multi-locus mixed model framework for the detection of small and linked QTLs in F2. Brief Bioinform 20(5), 1913–1924. 10.1093/bib/bby058.

Xing, Y., Tan, Y., Hua, J., Sun, X., Xu, C., Zhang, Q., 2002. Characterization of the main effects, epistatic effects and their environmental interactions of QTLs on the genetic basis of yield traits in rice. Theor Appl Genet 105(2-3), 248–257. 10.1007/s00122-002-0952-y.

Xu, S., 2013. Mapping quantitative trait Loci by controlling polygenic background effects. Genetics 195(4), 1209. 10.1534/genetics.113.157032.

Xu, S., Xu, Y., Gong, L., Zhang, Q., 2016. Metabolomic prediction of yield in hybrid rice. Plant J 88(2), 219–227. 10.1111/tpj.13242.

Xue, W., Xing, Y., Weng, X., Yu, Z., Tang, W., Lei, W., Zhou, H., Yu, S., Xu, C., Li, X., 2009. Natural variation in Ghd7 is an important regulator of heading date and yield potential in rice. Nat Genet 40(6), 761–767. 10.1038/ng.143.

Zaman, M.R., Paul, D.N.R., Akhter, N., Howlader, M.H., Kabir, M.S., 2006. Chi-square mixture of Chi-square distributions. J Appl Sci 6(2), 243–246. 10.3923/jas.2006.243.246.

Zhao, F., Xu, S., 2012a. An expectation and maximization algorithm for estimating Q×E interaction effects. Theor Appl Genet 124(8), 1375–1387. 10.1007/s00122-012-1794-x.

Zhao, F., Xu, S., 2012b. Genotype by environment interaction of quantitative traits: A case study in barley. G3-Genes Genom Genet 2(7), 779–788. 10.1534/g3.112.002980.

Zhao, F., Zhang, P., Wang, X., Akdemir, D., Garrick, D., He, J., Wang, L., 2023. Genetic gain and inbreeding from simulation of different genomic mating schemes for pig improvement. J Anim Sci Biotechnol 14(1), 87. 10.1186/s40104-023-00872-x.

Zhou, X., Stephens, M., 2012. Genome-wide efficient mixed model analysis for association studies. Nat Genet 44(7), 821–824. 10.1038/ng.2310.

Zhou, Y., Li, G., Zhang, Y., 2022. A compressed variance component mixed model framework for detecting small and linked QTL-by-environment interactions. Brief Bioinform 23(2), bbab596. 10.1093/bib/bbab596.

