## Supplementary material for "Identification of QTL-by-Environment Interaction by Controlling Polygenic Background Effect": File S4.docx

**Supplementary File S4** Variance components estimated for four traits from the 210 recombinant inbred lines of rice

| Trait | 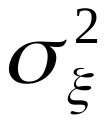 | 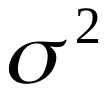 | 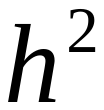 |
| --- | --- | --- | --- |
| KGW | 6.032 | 1.140 | 0.841 |
| TILLER | 1.0632 | 1.4551 | 0.4222 |
| GRAIN | 338.60 | 119.88 | 0.7395 |
| YIELD | 14.4028 | 19.7792 | 0.42135 |
