## Supplementary material for "Identification of QTL-by-Environment Interaction by Controlling Polygenic Background Effect": File S5.docx

Supplementary File S5. A example to demonstrate the new mixed model

The model is too complicated to understand. Let us use a small example to demonstrate the elements of the model in detail. Assume that
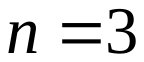
,
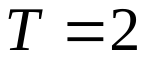
 and
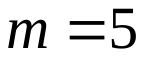
. First, let us consider an RIL population with two possible genotypes per locus. The following **Table S1** (top portion) shows the genotypes of the three individuals. Let us arbitrarily code one genotype as 0 and the other as 1. The numerical codes of genotypes are shown in the bottom portion of **Table S1**.

**Table** S1 Genotypes and their numerical codes of five markers for three individuals.

|  | Individual | marker1 | marker2 | marker3 | marker4 | marker5 |
| --- | --- | --- | --- | --- | --- | --- |
|  | 1 | AA | GG | AA | GG | CC |
| Genotype | 2 | AA | AA | CC | TT | TT |
|  | 3 | TT | GG | AA | TT | CC |
|  | 1 | 1 | 1 | 0 | 1 | 1 |
| Code | 2 | 1 | 0 | 1 | 0 | 0 |
|  | 3 | 0 | 1 | 0 | 0 | 1 |

Let us place the codes in an
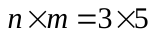
 matrix *Z*,

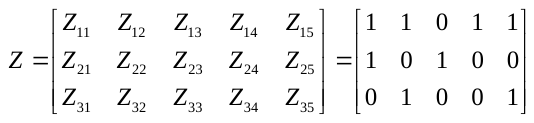
(S1)

Let us also define a marker inferred kinship matrix by

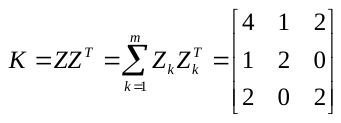
(S2)

where
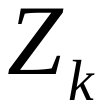
 is the *k*th column of matrix *Z*. For example, if the first marker is the current one under study, then
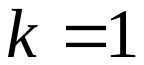
 and

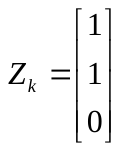

In general, the model can be written in the following form,

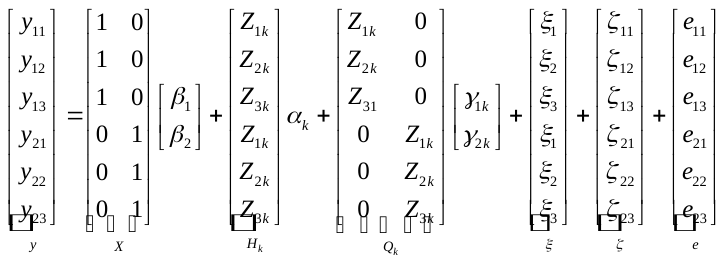
(S3)

where
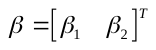
 and
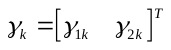
. Comparison of this model with model (1) will help understand the G×E model. The interaction effects (
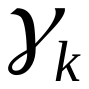
) can be treated either as fixed effects (if the number of environments is small) or random effects (if the number of environments is large). If treated as random effects, the G×E interaction effects should be assumed to have a
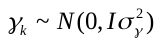
 distribution, where
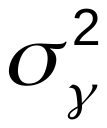
 is the interaction variance. In this study, we treat
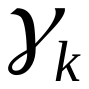
 as fixed effects. We also assume that
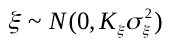
 and
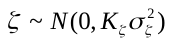
, where
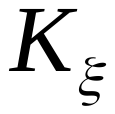
 is the kinship matrix for the polygenic main effects and
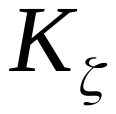
 be the kinship matrix for the polygenic G×E interaction effects. In this small example,

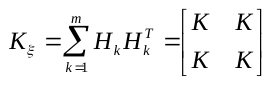
(S4)

and

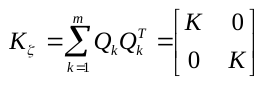
(S5)

If we assume that the residual error variance varies across different environments, then we should have two residual variances for
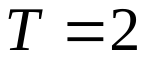
, which are denoted by
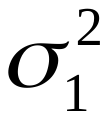
 and
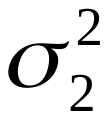
 for the two environments. Let us define two
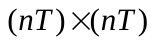
 matrices,

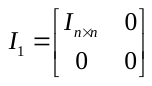
 and
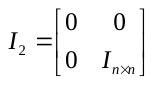

Let us treat the G×E interaction effects as fixed. We cannot estimate all levels of the interaction effects because the sum of all columns of matrix *Q_k_* is the same as matrix *H_k_*. However, this does not affect the significance test. We can examine the expectation and variance of data *y*, which are

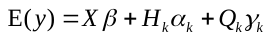
 (S6)

and

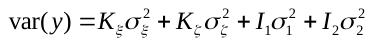
 (S7)

respectively. Detail of the expectation for
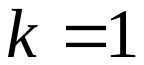
 is

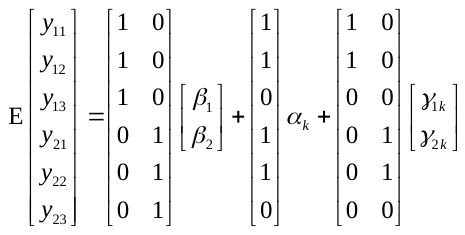
 (S8)

Components of the variance matrix are

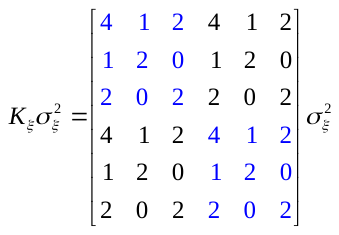

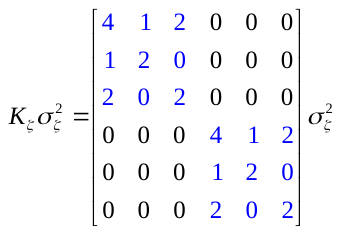

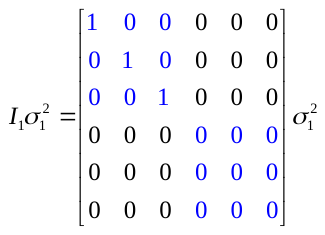

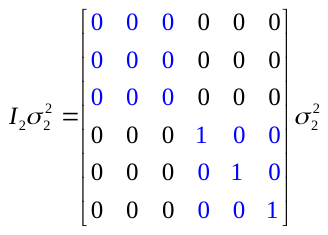

If we assume that the two environments have the same residual error variance, the variance should be

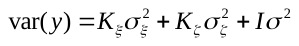
 (S9)

where

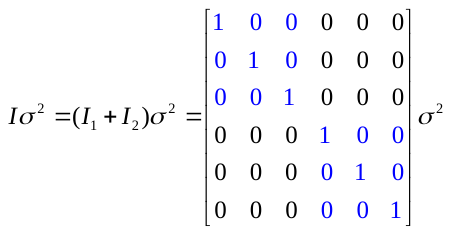

In this study, we assume that the residual error variance is homogenous across different environments and thus the variance matrix is
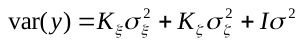
 and the expectation of *y* is
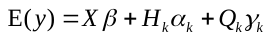
. In real data analysis, it is recommended to normalize the kinship matrix by dividing the original kinship matrix by the average of the diagonal elements of the original kinship matrix, i.e.,

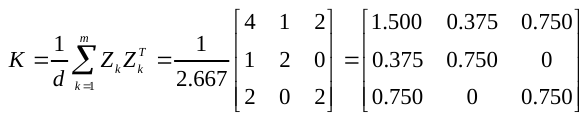
(S10)

where *d* is a normalization factor defined as

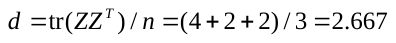
(S11)

Although normalization is optional, the estimated polygenic variances using the normalized kinship matrix can be compared with the residual error variance to give investigators a general idea on how large the polygenic variances are.
