## Supplementary material for "Identification of QTL-by-Environment Interaction by Controlling Polygenic Background Effect": File S6-S7.docx

**Supplementary** **File** **S6**: SAS PROC MIXED for Polygenic Variance Components Analysis

/*begin code*/

%let dir=D:\glm\sim;

filename bb "&dir\phe.csv";

filename kk1 "&dir\R1.csv";

filename kk2 "&dir\R2.csv";

filename gg "&dir\geno.csv";

**proc** **import** datafile=bb out=phe dbms=csv replace;

**proc** **import** datafile=kk1 out=kk1 dbms=csv replace;

**proc** **import** datafile=kk2 out=kk2 dbms=csv replace;

**proc** **import** datafile=gg out=gen dbms=csv replace;

**run**;

**proc** **mixed** data=phe method=ML nobound noprofile;

class year ril;

model yield = year/noint solution;

random ril/type=lin(**1**) ldata=kk1;

random ril/type=lin(**1**) ldata=kk2;

**run**;

/*end code*/

**Comments:** The program takes three input files stored in a user defined folder (D:\glm in this example), two files for the kinship matrices including polygenic main and interaction effects (named R1.csv and R2.csv in this example), one for the phenotypic values (named phe.csv in this example) and the fourth file for the bin genotypes (named geno.csv in this example). The Data R1.csv and R2.csv files must contain a variable for the id number of lines (named line in this example) and a variable for the phenotypic values of the trait in question (named yield in this example). The four input files are provided in Supplemental Datas S1, S2, S3 and S4, respectively.

Supplementary File S7: SAS PROC GLIMMIX for mapping main and Q×E interaction effects

/*begin code*/

%let dir=D:\glm;

filename bb "&dir\phe.csv";

filename kk1 "&dir\R1.csv";

filename kk2 "&dir\R2.csv";

filename gg "&dir\geno.csv";

**proc** **import** datafile=bb out=phe dbms=csv replace;

**proc** **import** datafile=kk1 out=kk1 dbms=csv replace;

**proc** **import** datafile=kk2 out=kk2 dbms=csv replace;

**proc** **import** datafile=gg out=gen dbms=csv replace;

**run**;

**proc** **iml**;

use gen;

read all into z;

close;

n=ncol(z);

m=nrow(z);

free zz;

do k=**1** to **1619**;

zk=z[k,]||z[k,]||z[k,]||z[k,];

zz=zz//zk`;

end;

create zz from zz;

append from zz;

close;

**quit**;

**data** one;

set phe;

do k=**1** to **1619**;

output;

end;

**run**;

**proc** **sort** data=one;

by k;

**run**;

**data** two;

merge one zz;

**run**;

**proc** **sort** data=two;

by k;

**run**;

**proc** **export** data=two outfile="gxedatatiller.csv" dbms=csv replace;

**run**;

**proc** **glimmix** data=two;

by k;

class year ril;

model tiller = year col1 col1*year/noint solution;

random ril /type=lin(**1**) ldata=kk1;

random ril /type=lin(**1**) ldata=kk2;

parms (**2.0769**) (**0.1171**) (**1.4581**) /hold=**1**,**2**;

ods output tests3=type3 ;

**run**;

**data** Main;

set type3;

if effect='COL1';

**run**;

**data** GxE;

set type3;

if effect='COL1*Year';

**run**;

**proc** **export** data=type3 outfile="D:\glm\tillerlastType3.csv" DBMS=csv replace;

**run**;

/*end code*/

**Comments:** The program takes three input files stored in a user defined folder (D:\glm\ in this example), one file for the kinship matrices including polygenic main and interaction effects (named R1.csv and R2.csv in this example), one for the phenotypic values (named phen99.csv in this example) and the third file for the bin genotypes (named geno.csv in this example). The Data R2.csv and R2.csv files must contain a variable for the id number of lines (named line in this example) and a variable for the phenotypic values of the trait in question (named kgw in this example). The four input files are provided in Supplemental Datas S1, S2, S3 and S4, respectively.
