## Supplementary figures and images for "Identification of QTL-by-Environment Interaction by Controlling Polygenic Background Effect"

### File S3.jpg

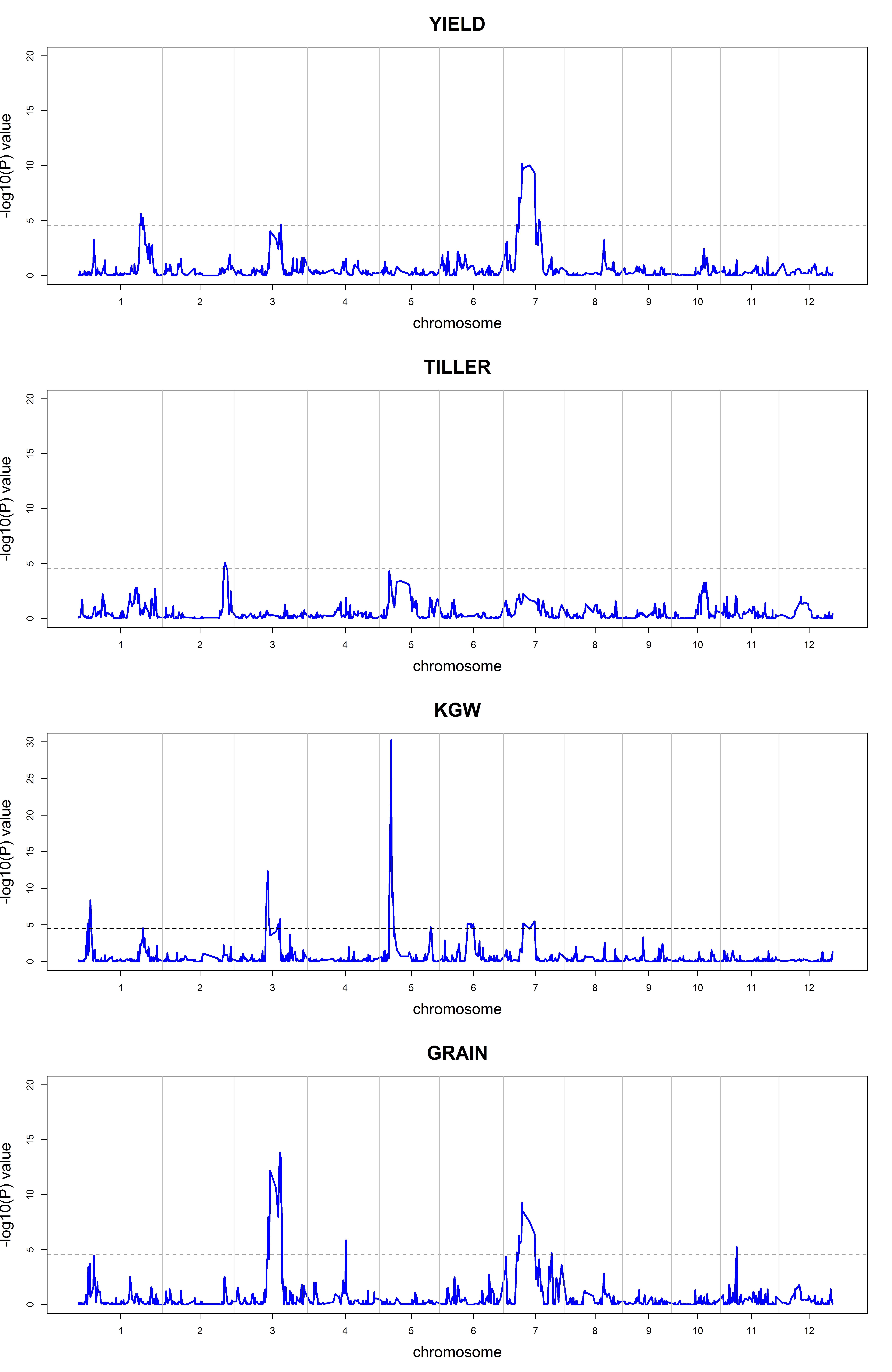
